# In vitro characterization of multidrug-resistant influenza A(H1N1)pdm09 viruses carrying a dual amino acid substitution associated with reduced susceptibility to neuraminidase inhibitors

**DOI:** 10.1101/334185

**Authors:** Emi Takashita, Seiichiro Fujisaki, Masaru Yokoyama, Masayuki Shirakura, Kazuya Nakamura, Tomoko Kuwahara, Noriko Kishida, Hironori Sato, Ikuko Doi, Yuji Sato, Shinichi Takao, Yukie Shimazu, Takeshi Shimomura, Takuo Ito, Shinji Watanabe, Takato Odagiri, The Influenza Virus Surveillance Group of Japan

## Abstract

We detected influenza A(H1N1)pdm09 viruses carrying dual H275Y/I223R, H275Y/I223K, or H275Y/G147R substitutions in their neuraminidase protein, respectively. These viruses showed cross-resistance to oseltamivir and peramivir and reduced susceptibility to zanamivir. The H275Y/G147R virus retained its replication capability at least in vitro, but the H275Y/I223R and H275Y/I223K viruses did not.

## Text

In Japan, four neuraminidase (NA) inhibitors––oseltamivir, peramivir, zanamivir, and laninamivir––are approved for the treatment of influenza. In addition, favipiravir, a viral RNA-dependent RNA polymerase inhibitor, was approved and stockpiled for use against novel influenza virus infections where existing antivirals are ineffective (1). The novel cap-dependent endonuclease inhibitor baloxavir marboxil was approved on 23 February 2018 for the treatment of influenza A and B virus infections and became available in hospitals from 14 March 2018 in Japan. Since nationwide monitoring is important for public health planning and clinical management, we have been conducting surveillance of antiviral-resistant viruses. In the 2013–2014 and 2015–2016 influenza seasons, we reported A(H1N1)pdm09 viruses exhibiting enhanced cross-resistance to oseltamivir and peramivir (2-4). These viruses possessed an I223R or a G147R substitution in combination with an H275Y substitution (N1 numbering) in their NA protein. In March 2016, we detected another dual H275Y mutant virus carrying an additional I223K substitution in its NA protein.

A few dual H275Y mutant viruses have been detected in immunocompromised and immunocompetent patients (5-7). Several studies have been carried out to understand the impact of the H275Y/I223R substitution on viral fitness (8, 9); however, that of the H275Y/I223K and H275Y/G147R viruses remains unknown. Here, we report our assessment of the in vitro properties of the dual H275Y mutant viruses isolated from immunocompromised patients.

First, we determined the NA inhibitor susceptibility of the dual H275Y mutant viruses by using a fluorescence-based NA assay with the NA-Fluor influenza neuraminidase assay kit (Applied Biosystems, Foster City, CA, USA). Oseltamivir carboxylate, peramivir, and zanamivir were purchased from Carbosynth Ltd. (Berkshire, United Kingdom). Laninamivir was kindly provided by Daiichi Sankyo Co. Ltd. (Tokyo, Japan). The H275Y/I223R, H275Y/I223K, and H275Y/G147R viruses exhibited cross-resistance to oseltamivir and peramivir and reduced susceptibility to zanamivir compared to the single H275Y viruses (Table 1) (3, 4). The H275Y/I223R and H275Y/I223K viruses, but not the H275Y/G147R virus, showed reduced susceptibility to laninamivir.

**TABLE 1.**
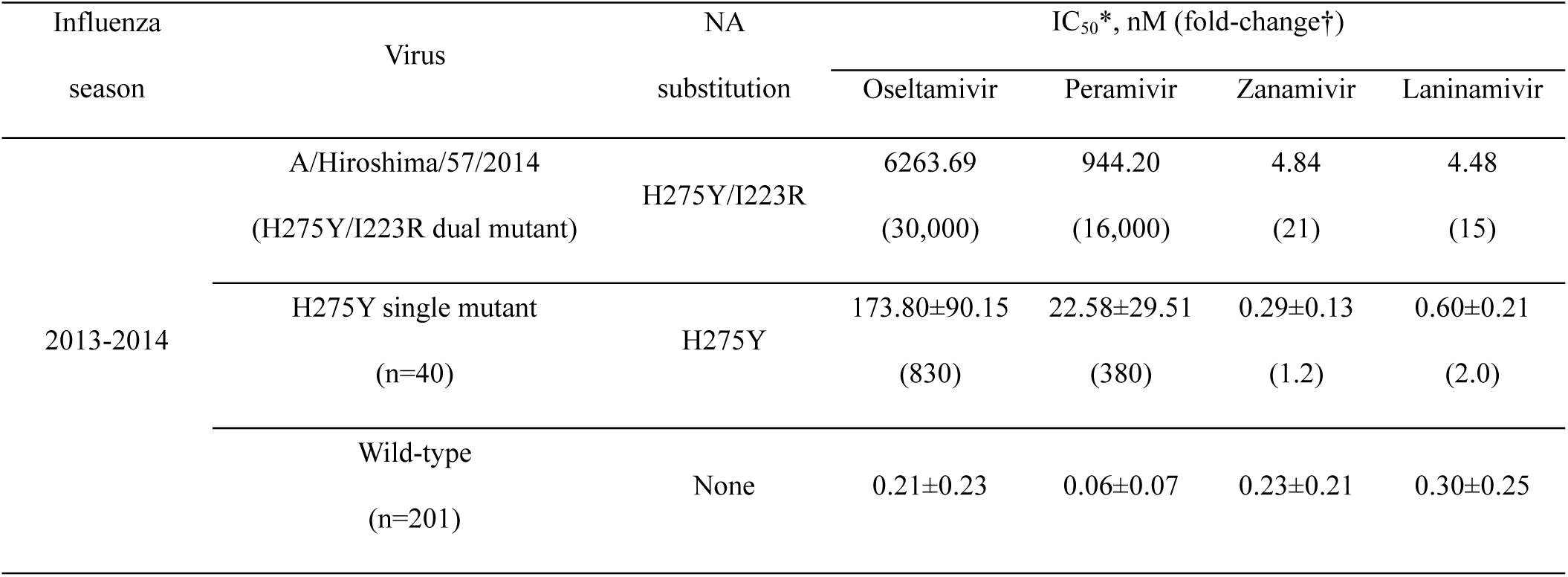
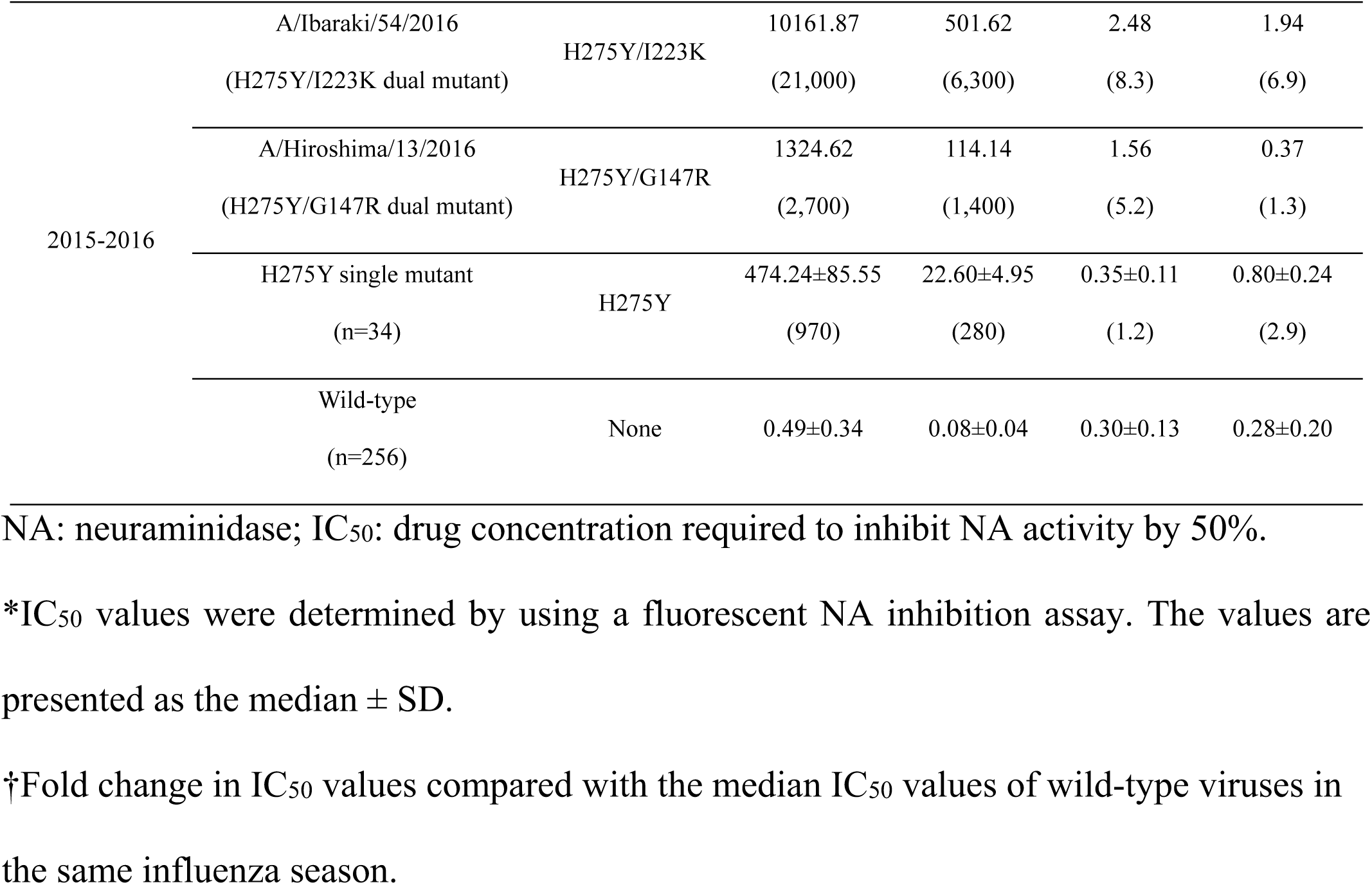
Susceptibility to neuraminidase inhibitors of H275Y dual mutant influenza A(H1N1)pdm09 viruses detected in Japan

We then analyzed representative single H275Y and wild-type viruses from the same genetic clade of each dual mutant virus in the same season (Table 2). The single H275Y and wild-type viruses possessed almost the same gene sequences as the dual mutant viruses except the dual substitutions in the NA. Statistical analyses were performed using GraphPad Prism version 6.0 for Mac OS X (GraphPad Software, La Jolla, CA, US). Statistically significant differences between groups were determined by using the Student’s t-test or Welch’s t-test on the result of the F-test. P values of <0.05 were considered statistically significant. All viruses tested were susceptible to favipiravir and no significant differences in susceptibility were found among the viruses (Table 2). Favipiravir was provided by Toyama Chemical Co. Ltd (Toyama, Japan).

**TABLE 2.**
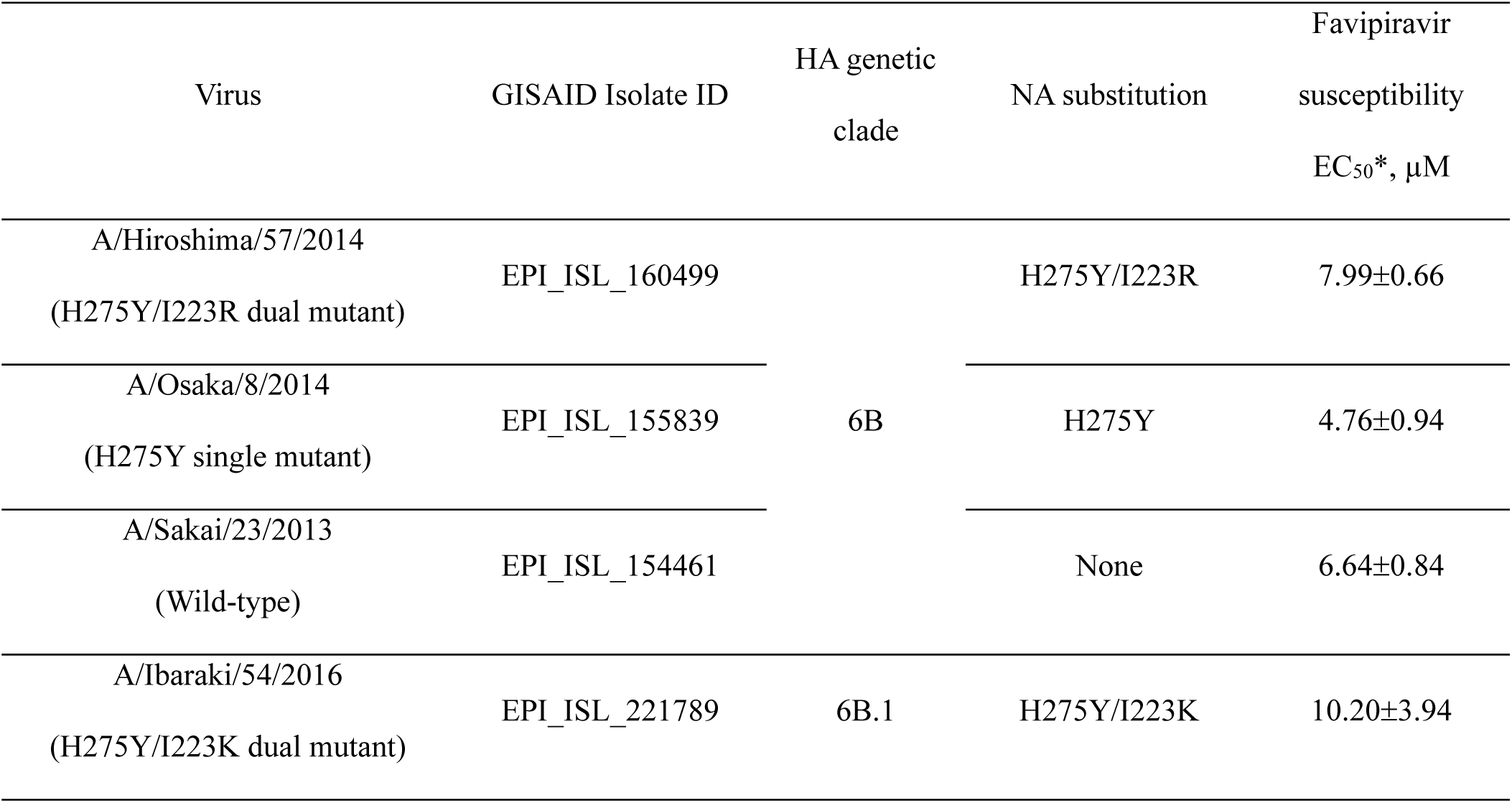
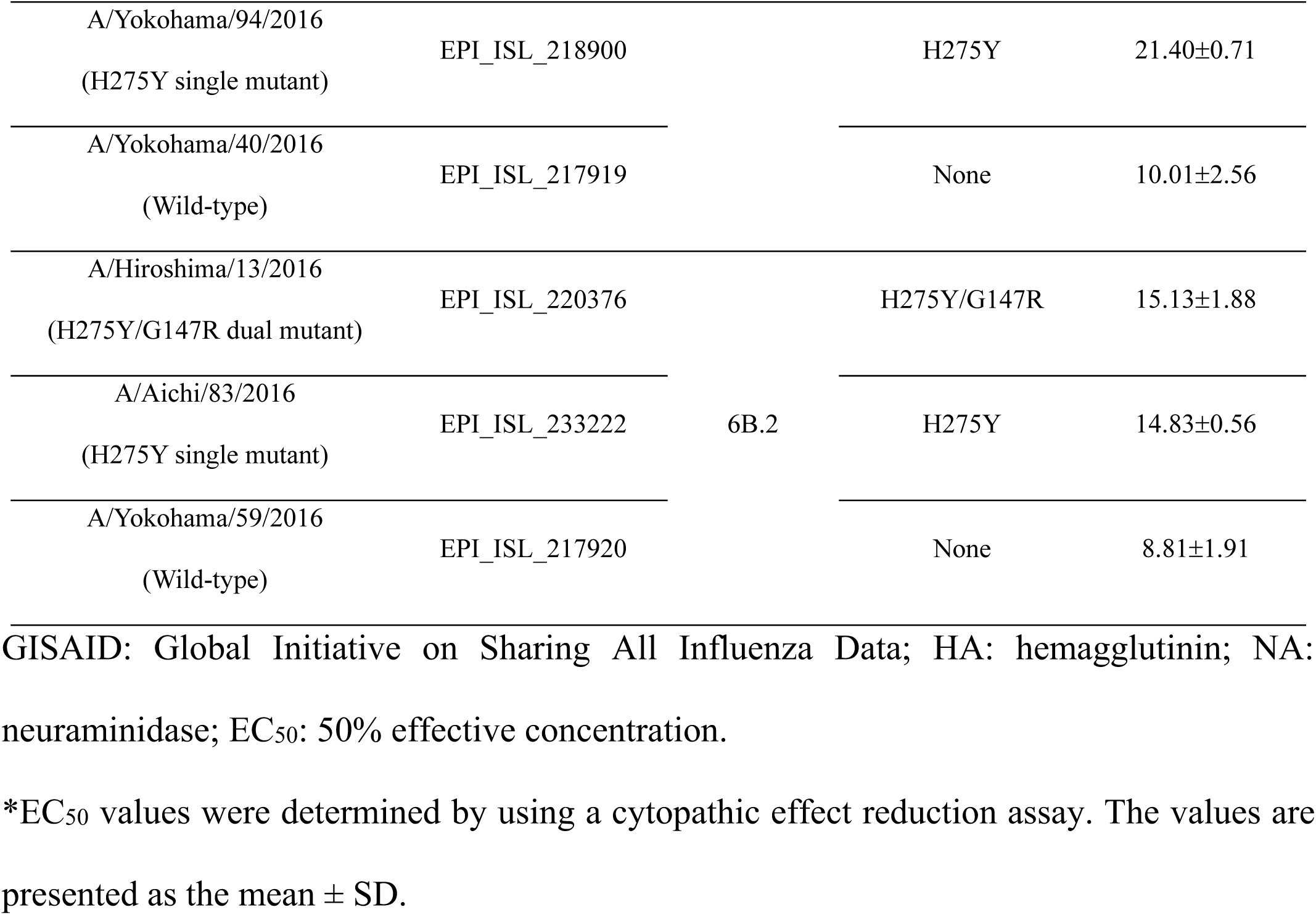
Influenza A(H1N1)pdm09 viruses used in this study

The NA activity of the H275Y/I223R and H275Y/I223K viruses was reduced compared with that of wild-type virus, consistent with a previous study (Figure 1A–C) (8). The H275Y/G147R virus showed comparable NA activity to that of the wild-type virus. These results suggest that the H275Y/I223R and H275Y/I223K substitutions are associated with a reduction in NA activity but that the H275Y/G147R substitution is not.

**FIG 1.**
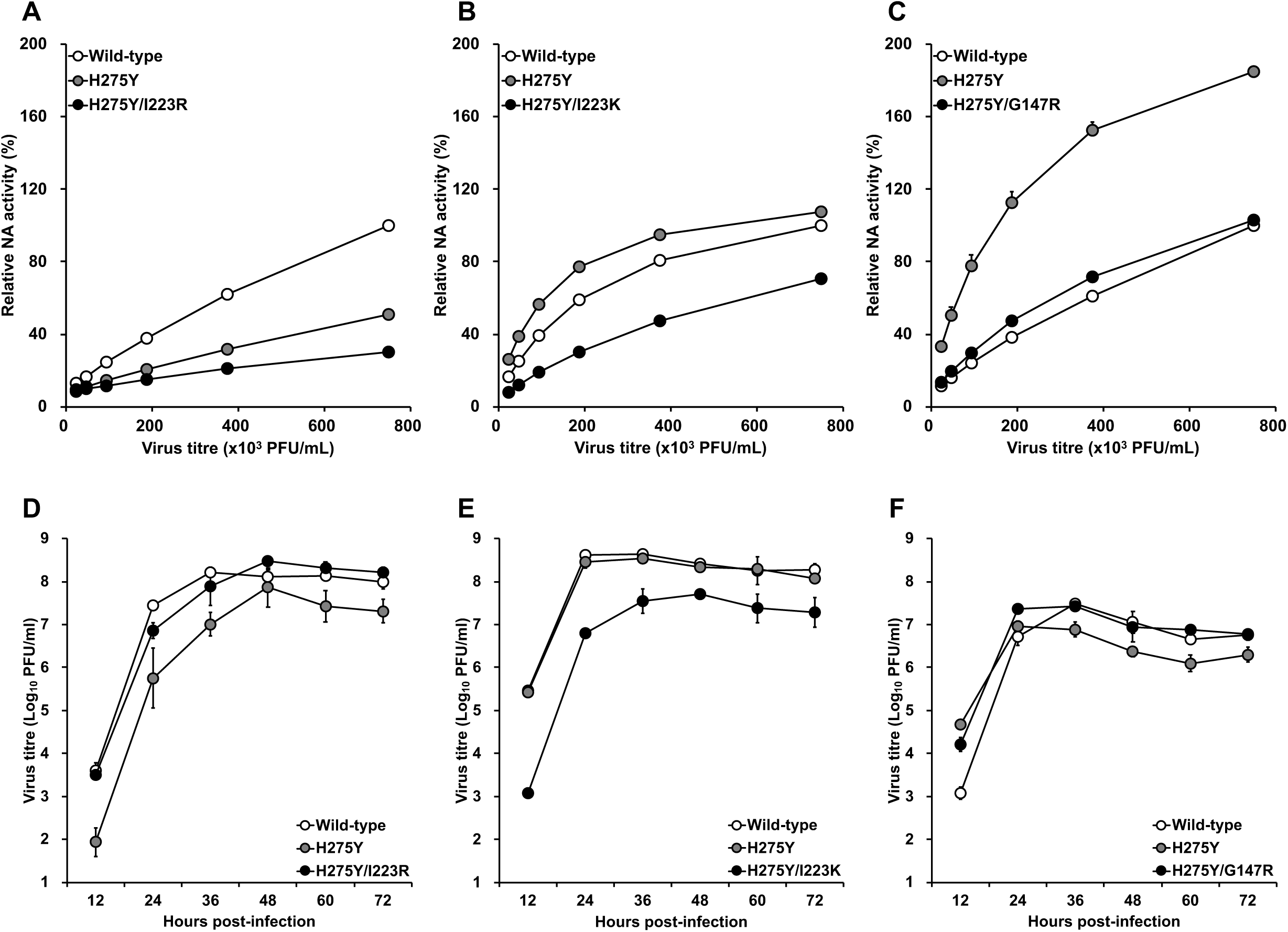
(A–C) Neuraminidase (NA) activities of the dual H275Y mutant influenza A(H1N1)pdm09 viruses. Serial 2-fold dilutions of viruses beginning at infectious doses of 7.5×10^5^ PFU/mL were subjected to the fluorescence-based NA assay in triplicate. The relative NA activities, normalized to that of the wild-type virus, are shown. Means and standard deviations are shown. (D–F) In vitro replication kinetics of the dual H275Y mutant influenza A(H1N1)pdm09 viruses. Confluent monolayers of MDCK-AX4 cells were infected in triplicate with viruses at a multiplicity of infection of 0.001 PFU/cell. The culture fluids were harvested at the indicated time points and were subjected to virus titration by using plaque assays in MDCK-AX4 cells. Means and standard deviations are shown. Wild-type: A/Sakai/23/2013 (A, D), A/Yokohama/40/2016 (B, E), or A/Yokohama/59/2016 (C, F); H275Y: A/Osaka/8/2014 (A, D), A/Yokohama/94/2016 (B, E), or A/Aichi/83/2016 (C, F).

The impact of the dual substitutions on viral growth was assessed using MDCK-AX4 cells (10), which overexpress the β-galactoside α2,6-sialyltransferase I gene (Figure 1D–F). MDCK-AX4 cells were kindly provided by Yoshihiro Kawaoka (University of Wisconsin, Madison, WI, US). Viral titres of the H275Y/I223R virus were comparable to those of the wild-type virus as previously described (9). The replication of the H275Y/I223K virus was significantly reduced compared with that of the single H275Y and the wild-type viruses. The H275Y/G147R and the wild-type viruses had comparable viral titres after 36 h post-infection, although the dual mutant virus replicated more efficiently than the wild-type virus during the initial cycle of infection. These results suggest that the H275Y/I223K substitutions negatively affected viral growth in vitro but that the H275Y/I223R and H275Y/G147R substitutions did not.

The competitive growth capability of each single or dual H275Y mutant virus with that of the wild-type virus was compared as previously described (Figure 2) (11). The proportion of the H275Y/I223R (Figure 2D) and H275Y/I223K (Figure 2E) viruses and their corresponding single H275Y viruses (Figures 2A and 2B) to that of wild-type viruses decreased significantly. However, the proportion of the single H275Y virus corresponding to the H275Y/G147R virus was comparable to that of wild-type virus (Figure 2C). Furthermore, the H275Y/G147R virus rapidly became dominant in the mixed virus populations at passages 1 and 2 (Figure 2F). These results indicate that the H275Y/G147R virus retained comparable growth ability to that of the wild-type virus, at least in vitro.

**FIG 2.**
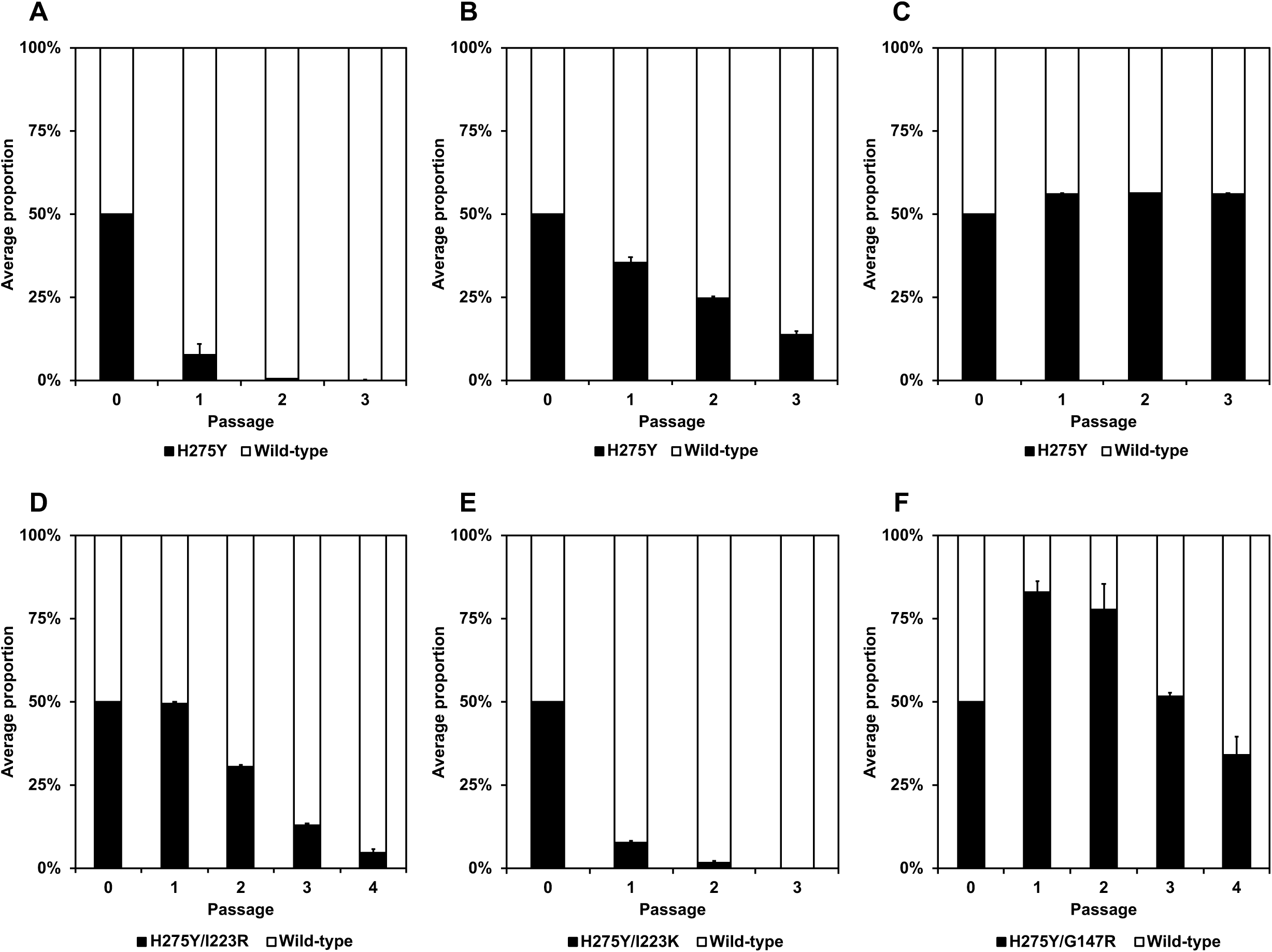
Competitive growth capabilities of the dual H275Y mutant influenza A(H1N1)pdm09 and wild-type viruses. MDCK-AX4 cells were coinfected in triplicate with the single H275Y (A–C), dual H275Y/I223R, H275Y/I223K, or H275Y/G147R (D–F) mutant virus and with the corresponding wild-type virus at a multiplicity of infection of 0.01 PFU/cell. At 2 days post-infection, the culture fluid was subjected to virus titration by using plaque assays in MDCK-AX4 cells and to deep sequencing analysis to determine the relative proportion of each genotype. The viruses were serially passaged 3–4 times at a multiplicity of infection of 0.01 PFU/cell. The error bars indicate the standard deviations. Wild-type: A/Sakai/23/2013 (A, D), A/Yokohama/40/2016 (B, E), or A/Yokohama/59/2016 (C, F); H275Y: A/Osaka/8/2014 (A), A/Yokohama/94/2016 (B), or A/Aichi/83/2016 (C).

To assess the potential effect of the amino acid substitutions on the stability of the NA, we performed an in silico mutagenesis study as previously described (11). The changes in stability caused by each of the substitutions I223R, I223K, and G147R were 1.32, 3.29, and −2.72 kcal/mol, respectively. The I223R and I223K substitutions were predicted to destabilise the NA structure, whereas the G147R substitution was predicted to stabilise the NA, which suggests that the G147R substitution compensates for structural disadvantages caused by the H275Y substitution (12).

The emergence of multidrug-resistant variants in patients treated with antiviral agents is a concern. The dual H275Y/I223R, H275Y/I223K, and H275Y/G147R viruses exhibited cross-resistance to oseltamivir and peramivir and reduced susceptibility to zanamivir. The patient who was infected with the H275Y/I223R virus recovered after laninamivir treatment; however, the other two died despite treatment with oseltamivir, peramivir and/or laninamivir. The dual H275Y mutant viruses were susceptible to favipiravir, suggesting that favipiravir could be a treatment option against these multidrug-resistant viruses.

The NA enzymatic active site includes catalytic residues that interact directly with the sialic acid substrate and framework residues that stabilize the active site (13). Residue 223 is located within the framework, and different substitutions at this position have different effects on NA inhibitor susceptibility (14). Residue 147 is located in a 150-loop, which are adjacent to the NA active site (2, 3). The G147R substitution may alter the conformation of the 150-loop due to the larger size and positive charge of the side chain, negatively affecting the binding of NA inhibitors (2, 3). In this study, the NA activity of the H275Y/I223R and H275Y/I223K viruses was significantly reduced, whereas the H275Y/G147R virus retained its NA activity. These differences may be explained by the positions of amino acids, inside or outside, the NA active site.

The single H275Y virus corresponding to the H275Y/G147R virus showed significantly increased NA activity, but reduced growth capability compared to the wild-type virus. However, the NA activity and growth capability of the H275Y/G147R virus was comparable to that of the wild-type virus. An optimal balance between hemagglutinin (HA) receptor-binding and NA enzymatic activities is important for growth and transmissibility of influenza viruses (15-17). Mismatched HA and NA pairs can be rescued by amino acid substitutions that compensate for increased or decreased activities (18). Thus, the introduction of the G147R substitution into the H275Y mutant NA may compensate for the high NA activity, resulting in the restoration of viral growth.

Our structural analysis predicts that the I223R and I223K substitutions destabilise the NA structure, but the stability score for I223R was lower than that for I223K. These results suggest that an NA with the I223R substitution is more stable than an NA with the I223K substitution.

Hooper et al. reported that the G147R substitution confers receptor-binding activity to the NA proteins and moderate resistance to neutralization by the Fab of a monoclonal antibody against the HA receptor-binding pocket (19). In this study, we showed that the H275Y/G147R virus retained its replication capability at least in vitro. In fact, the patient infected with the H275Y/G147R virus developed pneumonia without isolation of bacterial pathogens, suggesting viral pneumonia with this dual mutant virus (3).

The multidrug-resistant viruses carrying the dual H275Y substitution were detected in immunocompromised patients after prolonged treatment with one or more NA inhibitors. Therefore, the emergence of resistant viruses during NA inhibitor administration should be closely monitored to improve clinical management. Furthermore, the surveillance of antiviral-resistant viruses should be continued to protect public health.

## Acknowledgments

We thank Rie Ogawa, Hideka Miura, Miki Akimoto, Aya Sato, Kayo Watanabe, and Hiromi Sugawara for technical assistance. We also thank Susan Watson for scientific editing.

This work was supported by a Grant-in-Aid for Emerging and Reemerging Infectious Diseases from the Ministry of Health, Labour and Welfare, Japan (grant no. 10110400) and by JSPS KAKENHI Grant number 18K10036.

Members of the Influenza Virus Surveillance Group of Japan are Rika Komagome (Hokkaido Institute of Public Health), Asami Ohnishi (Sapporo City Institute of Public Health), Rika Tsutsui (Aomori Prefectural Public Health and Environment Center), Masaki Takahashi (Iwate Prefectural Research Institute for Environmental Sciences and Public Health), Yuko Suzuki (Miyagi Prefectural Institute of Public Health and Environment), Makiko Ushimizu (Sendai City Institute of Public Health), Chihiro Shibata (Akita Prefectural Research Center for Public Health and Environment), Shizuka Tanaka (Yamagata Prefectural Institute of Public Health), Yoshiko Kashiwagi (Fukushima Prefectural Institute of Public Health), Chika Hirokawa (Niigata Prefectural Institute of Public Health and Environmental Sciences), Kazunari Yamamoto (Niigata City Institute of Public Health and Environment), Takako Suzuki (Tochigi Prefectural Institute of Public Health and Environmental Sciences), Shunsuke Kataoka (Utsunomiya City Institute of Public Health and Environment Science), Hiroyuki Tsukagoshi (Gunma Prefectural Institute of Public Health and Environmental Sciences), Noriko Suzuki (Saitama Institute of Public Health), Yuka Uno (Saitama City Institute of Health Science and Research), Noriko Oitate (Chiba Prefectural Institute of Public Health), Wakako Nishikawa (Chiba City Institute of Health and Environment), Sachiko Harada (Tokyo Metropolitan Institute of Public Health), Sumi Watanabe (Kanagawa Prefectural Institute of Public Health), Chiharu Kawakami (Yokohama City Institute of Public Health), Hideaki Shimizu (Kawasaki City Institute of Public Health), Hazime Amano (Yokosuka Institute of Public Health), Sayoko Arakawa (Sagamihara City Institute of Public Health), Masayuki Oonuma (Yamanashi Institute for Public Health), Michiko Takeuchi (Nagano Environmental Conservation Research Institute), Yuichiro Okamura (Nagano City Health Center), Yukiko Sakai (Shizuoka Institute of Environment and Hygiene), Takaharu Maehata (Shizuoka City Institute of Environmental Sciences and Public Health), Toshihiko Furuta (Hamamatsu City Health Environment Research Center), Masatsugu Obuchi (Toyama Institute of Health), Hiroe Kodama (Ishikawa Prefectural Institute of Public Health and Environmental science), Kaori Sato (Fukui Prefectural Institute of Public Health and Environmental Science), Masahiro Nishioka (Gifu Prefectural Research Institute for Health and Environmental Sciences), Yusuke Sato (Gifu Municipal Institute of Public Health), Yoshihiro Yasui (Aichi Prefectural Institute of Public Health), Takuya Yano (Mie Prefecture Health and Environment Research Institute), Hiromi Kodama (Shiga Prefectural Institute of Public Health), Akiko Nagasao (Kyoto City Institute of Health and Environmental Sciences), Satoshi Hiroi and Hideyuki Kubo (Osaka Institute of Public Health), Fumika Okayama (Sakai City Institute of Public Health), Tomohiro Oshibe (Hyogo Prefectural Institute of Public Health and Consumer Sciences), Ai Mori (Kobe Institute of Health), Misako Fujitani (Nara Prefecture Institute of Health), Yuki Matsui (Wakayama Prefectural Research Center of Environment and Public Health), Hidenobu Ekawa (Wakayama City Institute of Public Health), Nobuyuki Kato (Tottori Prefectural Institute of Public Health and Environmental Science), Tetsuo Mita (Shimane Prefectural Institute of Public Health and Environmental Science), Yasuhiro Matsuoka (Okayama Prefectural Institute for Environmental Science and Public Health), Miwako Yamamoto (Hiroshima City Institute of Public Health), Shoichi Toda (Yamaguchi Prefectural Institute of Public Health and Environment), Yumiko Kawakami (Tokushima Prefectural Public Health, Pharmaceutical and Environmental Sciences Center), Yukari Terajima (Kagawa Prefectural Research Institute for Environmental Sciences and Public Health), Akie Ochi (Ehime Prefecture Institute of Public Health and Environmental Science), Noriko Yorimitsu (Kochi Public Health and Sanitation Institute), Yuki Ashizuka (Fukuoka Institute of Health and Environmental Sciences), Shuichi Zaitsu (Fukuoka City Institute of Health and Environment), Takashi Kimura (Kitakyushu City Institute of Health and Environmental Sciences), Katsuyuki Ando (Saga Prefectural Institute of Public Health and Pharmaceutical Research), Kana Miura (Nagasaki Prefectural Institute for Environment Research and Public Health), Kenta Yoshioka (Kumamoto Prefectural Institute of Public-Health and Environmental Science), Kaori Nishizawa (Kumamoto City Environmental Research Center), Miki Kato (Oita Prefectural Institute of Health and Environment), Miho Miura (Miyazaki Prefectural Institute for Public Health and Environment), Yuka Iwamoto (Kagoshima Prefectural Institute for Environmental Research and Public Health), and Yumani Kuba (Okinawa Prefectural Institute of Health and Environment).

## References

1. Takashita E, Ejima M, Ogawa R, Fujisaki S, Neumann G, Furuta Y, Kawaoka Y, Tashiro M, Odagiri T. 2016. Antiviral susceptibility of influenza viruses isolated from patients pre- and post-administration of favipiravir. Antiviral Res 132:170–7.

2. Gubareva LV, Besselaar TG, Daniels RS, Fry A, Gregory V, Huang W, Hurt AC, Jorquera PA, Lackenby A, Leang SK, Lo J, Pereyaslov D, Rebelo-de-Andrade H, Siqueira MM, Takashita E, Odagiri T, Wang D, Zhang W, Meijer A. 2017. Global update on the susceptibility of human influenza viruses to neuraminidase inhibitors, 2015-2016. Antiviral Res 146:12–20.

3. Takashita E, Fujisaki S, Shirakura M, Nakamura K, Kishida N, Kuwahara T, Shimazu Y, Shimomura T, Watanabe S, Odagiri T. 2016. Influenza A(H1N1)pdm09 virus exhibiting enhanced cross-resistance to oseltamivir and peramivir due to a dual H275Y/G147R substitution, Japan, March 2016. Euro Surveill 21.

4. Takashita E, Meijer A, Lackenby A, Gubareva L, Rebelo-de-Andrade H, Besselaar T, Fry A, Gregory V, Leang SK, Huang W, Lo J, Pereyaslov D, Siqueira MM, Wang D, Mak GC, Zhang W, Daniels RS, Hurt AC, Tashiro M. 2015. Global update on the susceptibility of human influenza viruses to neuraminidase inhibitors, 2013-2014. Antiviral Res 117:27–38.

5. "Anonymous. 2009. Oseltamivir-resistant 2009 pandemic influenza A (H1N1) virus infection in two summer campers receiving prophylaxis--North Carolina, 2009. MMWR Morb Mortal Wkly Rep 58:969–72.

6. Nguyen HT, Fry AM, Loveless PA, Klimov AI, Gubareva LV. 2010. Recovery of a multidrug-resistant strain of pandemic influenza A 2009 (H1N1) virus carrying a dual H275Y/I223R mutation from a child after prolonged treatment with oseltamivir. Clin Infect Dis 51:983–4.

7. Nguyen HT, Trujillo AA, Sheu TG, Levine M, Mishin VP, Shaw M, Ades EW, Klimov AI, Fry AM, Gubareva LV. 2012. Analysis of influenza viruses from patients clinically suspected of infection with an oseltamivir resistant virus during the 2009 pandemic in the United States. Antiviral Res 93:381–6.

8. LeGoff J, Rousset D, Abou-Jaoude G, Scemla A, Ribaud P, Mercier-Delarue S, Caro V, Enouf V, Simon F, Molina JM, van der Werf S. 2012. I223R mutation in influenza A(H1N1)pdm09 neuraminidase confers reduced susceptibility to oseltamivir and zanamivir and enhanced resistance with H275Y. PLoS One 7:e37095.

9. Pizzorno A, Abed Y, Bouhy X, Beaulieu E, Mallett C, Russell R, Boivin G. 2012. Impact of mutations at residue I223 of the neuraminidase protein on the resistance profile, replication level, and virulence of the 2009 pandemic influenza virus. 9. Antimicrob Agents Chemother 56:1208–14.

10. Hatakeyama S, Sakai-Tagawa Y, Kiso M, Goto H, Kawakami C, Mitamura K, Sugaya N, Suzuki Y, Kawaoka Y. 2005. Enhanced expression of an alpha2,6-linked sialic acid on MDCK cells improves isolation of human influenza viruses and evaluation of their sensitivity to a neuraminidase inhibitor. J Clin Microbiol 43:4139–46.

11. Takashita E, Kiso M, Fujisaki S, Yokoyama M, Nakamura K, Shirakura M, Sato H, Odagiri T, Kawaoka Y, Tashiro M. 2015. Characterization of a large cluster of influenza A(H1N1)pdm09 viruses cross-resistant to oseltamivir and peramivir during the 2013-2014 influenza season in Japan. Antimicrob Agents Chemother 59:2607–17.

12. Hurt AC, Hardie K, Wilson NJ, Deng YM, Osbourn M, Leang SK, Lee RT, Iannello P, Gehrig N, Shaw R, Wark P, Caldwell N, Givney RC, Xue L, Maurer-Stroh S, Dwyer DE, Wang B, Smith DW, Levy A, Booy R, Dixit R, Merritt T, Kelso A, Dalton C, Durrheim D, Barr IG. 2012. Characteristics of a widespread community cluster of H275Y oseltamivir-resistant A(H1N1)pdm09 influenza in Australia. J Infect Dis 206:148–57.

13. Colman PM, Hoyne PA, Lawrence MC. 1993. Sequence and structure alignment of paramyxovirus hemagglutinin-neuraminidase with influenza virus neuraminidase. Journal of Virology 67:2972–2980.

14. Samson M, Pizzorno A, Abed Y, Boivin G. 2013. Influenza virus resistance to neuraminidase inhibitors. Antiviral Res 98:174–85.

15. Lakdawala SS, Lamirande EW, Suguitan AL, Jr., Wang W, Santos CP, Vogel L, Matsuoka Y, Lindsley WG, Jin H, Subbarao K. 2011. Eurasian-origin gene segments contribute to the transmissibility, aerosol release, and morphology of the 2009 pandemic H1N1 influenza virus. PLoS Pathog 7:e1002443.

16. Xu R, Zhu X, McBride R, Nycholat CM, Yu W, Paulson JC, Wilson IA. 2012. Functional balance of the hemagglutinin and neuraminidase activities accompanies the emergence of the 2009 H1N1 influenza pandemic. J Virol 86:9221–32.

17. Yen HL, Liang CH, Wu CY, Forrest HL, Ferguson A, Choy KT, Jones J, Wong DD, Cheung PP, Hsu CH, Li OT, Yuen KM, Chan RW, Poon LL, Chan MC, Nicholls JM, Krauss S, Wong CH, Guan Y, Webster RG, Webby RJ, Peiris M. 2011. Hemagglutinin-neuraminidase balance confers respiratory-droplet transmissibility of the pandemic H1N1 influenza virus in ferrets. Proc Natl Acad Sci U S A 108:14264–9.

18. Mitnaul LJ, Matrosovich MN, Castrucci MR, Tuzikov AB, Bovin NV, Kobasa D, Kawaoka Y. 2000. Balanced hemagglutinin and neuraminidase activities are critical for efficient replication of influenza A virus. J Virol 74:6015–20.

19. Hooper KA, Crowe JE, Jr., Bloom JD. 2015. Influenza viruses with receptor-binding N1 neuraminidases occur sporadically in several lineages and show no attenuation in cell culture or mice. J Virol 89:3737–45.

